# An Option Space for Early Neural Evolution

**DOI:** 10.1101/027425

**Authors:** Gáspár Jékely, Fred Keijzer, Peter Godfrey Smith

## Abstract

The origin of nervous systems has traditionally been discussed within two conceptual frameworks. Input-output models stress the sensory-motor aspects of nervous systems, while internal coordination models emphasize the role of nervous systems in coordinating multicellular activity, especially muscle-based motility. Here we consider both frameworks and apply them to describe aspects of each of three main groups of phenomena that nervous systems control: behavior, physiology and development. We argue that both frameworks and all three aspects of nervous system function need to be considered for a comprehensive discussion of nervous system origins. This broad mapping of the option space enables an overview of the many influences and constraints that may have played a role in the evolution of the first nervous systems.

## 1. Introduction

The origin of the nervous system was an evolutionary event that fundamentally changed how control is achieved within a multicellular body. Recent progress in genomics, phylogenetics, developmental biology and the study of simple nervous systems has provided a wealth of new empirical information that bears on the earliest stages in neural evolution. However, many of the conceptual frameworks that are used to discuss this work recognize only a limited subset of the range of roles that nervous systems can play; the neural control of development and physiology, in addition to behavior, is often sidelined or omitted. In addition, these frameworks tend to employ an overly simple conception of the role of neural activity in the adaptive shaping of behavior itself. The aim of this paper is to organize ideas and hypotheses in this area in a global way by charting the ‘option space’ for hypotheses about early neural evolution, making explicit the entire range of functions that early nervous systems may have played.

Historically, the origin of nervous systems has been discussed in light of two different conceptual models. We call these the *input-output* (IO) and *internal coordination* (IC) models. The two models emphasize two different aspects of the nervous system as a control device. According to IO models, the main role of the nervous system is to receive sensory information and process it to produce meaningful motor output. Braitenberg’s ‘vehicles’ [1] represent a simple conceptual IO model of an organism, where directional light sensors modify the speed of wheels in a moving vehicle.

In contrast to IO models, IC models hold that a central role of early nervous systems was to induce and coordinate activity internal to large multicellular organizations. While an IO model tends to assume an operational effector system and addresses how this system is to be put to use, an IC model highlights the evolutionary shift involved in generating new multicellular effectors. In particular, the use of extensive contractile tissues – muscle – by large organisms is an important evolutionary invention. Achieving organized movement in a muscle is a demanding task that should not be taken for granted, as sometimes happens in discussions employing an IO framework.

The difference between IC and IO models can be understood more abstractly as a distinction between two kinds of coordination. An IO device aims to coordinate what is done by the organism with the state of the environment; it is concerned with *act-state* coordination, where the difference between acts and states is that acts are choices of the organism itself while states are external and must be sensed.

The aim of an IC device, in contrast, is to coordinate different aspects of what an organism *does;* it is concerned with *act-act* coordination. Expressed differently, it coordinates the micro-acts of a system’s parts into the macro-acts of a whole (Figure 1). We argue that both these conceptual frameworks must be considered when discussing early nervous system evolution. The IO/IC distinction is applied here both to conceptual frameworks used to explain nervous system phenomena and also to ways a nervous system can actually be organized. We will refer to IO and IC ‘models’ and ‘systems’ to refer to conceptual frameworks and to types of nervous system organization, respectively.

**Figure 1.**
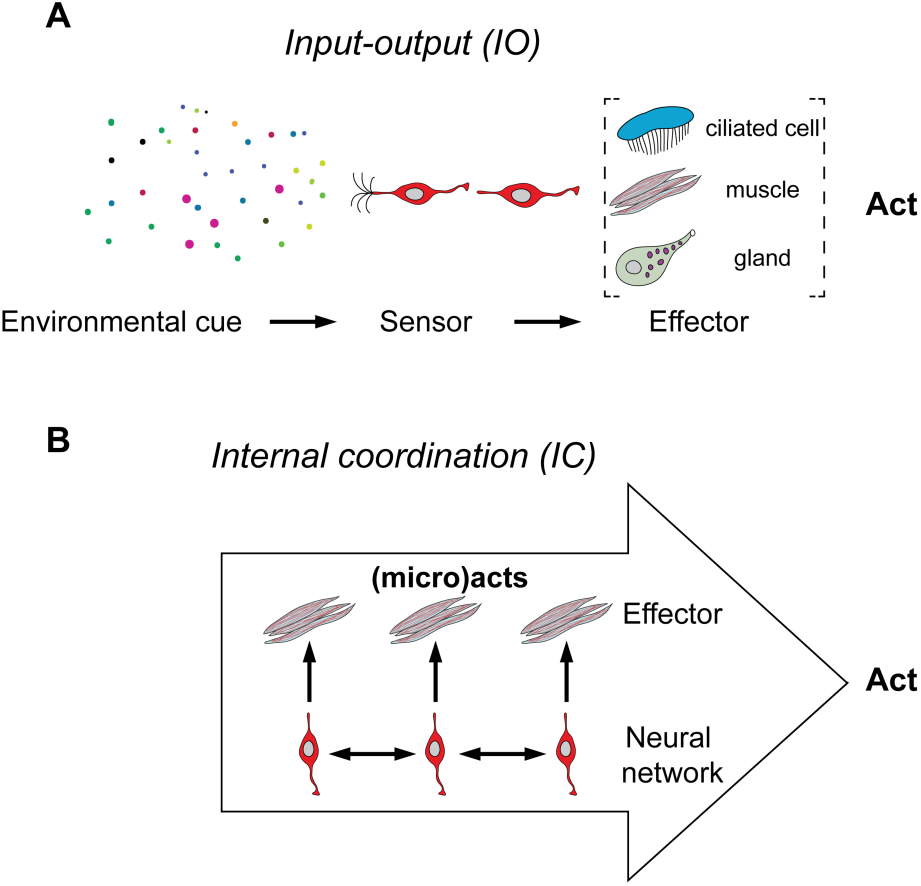
Schematic of input-output and internal coordination systems.

Historically, there has been a strong emphasis on IO models in attempts to understand nervous system function and early neural evolution. This mainstream tradition was heavily influenced by Charles Sherrington’s work on the reflex arc [2]. The reflex arc was used as a paradigm case of neural organization by G.H. Parker, who proposed an influential scenario for the origin of sensory-motor coordination. As Parker saw it, "independent effectors” arose first, and the essential function of early nervous systems was to connect these effectors with specialized sensors (“receptor mechanisms”, [3]). This tradition runs through to the present day and is seen especially in work on the evolution of locomotion systems, as exemplified by Mackie [4] [5] and Jékely [6].

IC models date from the 1950s, especially with the work of Chris Pantin [7], who criticized Parker for failing to recognize the importance of internal coordination in the new effector systems that arose in metazoan evolution, especially with the origin of muscle. As Pantin said, “the complex and important movements required in behaviour can only be brought about indirectly through the co-ordinated contraction of large regions of these muscle sheets…, which indirectly move and distort the body into the position or shape required.” [7]. IC models also stress the role of coordinators of endogenous activity such as oscillators, which initiate and maintain coordinated activity across muscle tissue that is not directly linked to sensory input [8]. While initially taken up by some biologists such as Passano [8] and de Ceccatty [9], the IC model never became very prominent, although recently it has been reintroduced by Keijzer, van Duijn and Lyon [10]. Our present aim is to combine the IO and IC models into a single framework; we argue that both need to be considered to understand early neural evolution as they stress different but complementary issues. In the sections to follow, we apply these two frameworks to describe aspects of each of three main groups of phenomena that nervous systems control: behavior, physiology and development. This broad mapping of the option space enables a more comprehensive overview of the many influences and constraints that played a role in the evolution of the first nervous systems.

## 2. What are nervous systems and what do they do?

A nervous system represents the totality of neurons in an organism. The definition of a *neuron* is more difficult than it initially appears, however. Many standard definitions are too narrow. For example, if the presence of synapses is required, then this results in the exclusion of some important activities even within our own brains, where neurosecretory cells influence other cells via paracrine or hormonal signaling. The most fundamental features of neural activity are excitability, and the influencing of the activity of other cells on small temporal scales (milliseconds to seconds). This can happen either by synaptic (chemical or electrical) or neuroendocrine (paracrine, hormonal) means. It would be possible to use a very broad and purely functional concept of “neuron,” in which a neuron is any electrically excitable cell that influences another cell by means of electrical or secretory mechanisms. This broad definition would include some cases that are not usually seen as neural phenomena – for example, electrically conducting tissues in some plants – and it might be unhelpfully broad for that reason. We suggest that it will be clearest to use a concept narrower than that one, but still broader than many textbook definitions. This definition augments the broad functional view of the neuron with an anatomical requirement – we only include excitable cells with specialized projections, such as axons and dendrites. A neuron in our sense, then, is *an electrically excitable cell that influences another cell by means of electrical or secretory mechanisms, and whose morphology includes specialized projections.* This, or any other, definition of “neuron” is best employed with an expectation that we will encounter grey areas and borderline cases. Here, as elsewhere in biology, it is important to avoid what Ernst Mayr called “typological thinking,” the imposition of sharp boundaries in domains where such boundaries are unlikely to be found.

Our option space for neural evolution distinguishes three roles that nervous systems play in an organism: the control of *behavior,* the control of *physiology,* and the control of *development.* We will argue that all three roles probably have considerable evolutionary importance. The control of behavior is the most familiar role, and workers in cognitive science tend to emphasize it above everything else when discussing nervous system function. The control of behavior includes phenomena like locomotor control, along with the control of sexual behavior and feeding. The control of physiology, a second major role of nervous systems, includes phenomena such as circadian and circa-lunar clocks, the control of metabolism, digestion, and diuresis. Some borderline cases can be categorized either as behavior or physiology, such as the feeding and peristaltic gut motion in a sea anemone.

The control of development, also neglected in many discussions, is a fundamental role of all animal nervous systems. It includes the control of growth and metamorphosis, along with phenomena such as molting and regeneration. These processes are controlled by hormonal signals emanating from the nervous system. The next three sections discuss each of these major families of functions in the light of IO and IC models.

## 3. Behavior

Animals make use of three basic kinds of effector systems for the production of behavior: ciliary motion, muscular contraction, and glandular secretion (Figure 1). This list is not exhaustive (the activity of bioluminescent photocytes in ctenophores, for example, is distinct from these), but it covers the main types.

Though the centrality of these effector systems is uncontroversial, the boundary between “behavior” and other phenomena may appear somewhat different from the vantage point of IO and IC models. An IO model tends to cast behavior in relation to environmental factors and functional environmental effects; an IC model stresses self-generated motion that imposes a force on some medium as the key feature of behavior, irrespective of whether or how this has specific environmental effects. Thus coordinating heart muscle in the pumping of blood, or coordinating lung cilia to expel mucus would be clear forms of behavior from an IC viewpoint, while remaining boundary cases in the former.

### 3.1 IC and IO systems for cilia-based behaviors

Ciliary beating is used for locomotion in a wide range of small organisms, and it has other uses as well; inside a sponge, for example, cilia are used to create water flow to enable access to food and oxygen. Many marine larvae employ cilia to bring food into their mouth [11–13]. In all these cases, the cilia must have their movements *coordinated* – this is a first context in which an IC function might be relevant. However, coordination of cilia can often be achieved by non-neural means. In particular, adjacent cilia in multiciliated epithelia spontaneously synchronize their beating activity by means of hydrodynamic coupling, leading to the formation of metachronal waves in multiciliated epithelia or ciliary bands [14–16]

The large-scale non-neural coordination of cilia requires that cilia themselves be properly oriented within the body, though. This is ensured by the planar polarity of the cells, which controls the axis of beating [17]. In ciliated larvae, Wnt signaling is the likely ancestral regulator of establishing axial polarity of the body and the planar polarity of cilia. In the ciliated larvae of the cnidarian *Clytia hemisphaerica,* a Wnt ligand is expressed in the oral pole [18] and directly or indirectly regulates PCP signaling [19]. The planar polarity of the ciliated epithelium requires the conserved protein strabismus both in cnidarians and vertebrates [19,20]. Sponge larvae also have polarized ciliated epithelia, express a Wnt in the posterior pole [21], and the sponge genome contains the conserved components of PCP signaling [22]. If the planar polarity of cilia is established, the coordinated beating of cilia emerges via physical principles. So in this case, there is signaling in *development* that sets things up so that the IC function in the cilia themselves can be achieved *without* signaling (or other internal control devices) during behavior.

Once coordinated ciliary motion exists in an organism, control devices may modify the activity of the cilia. Thus cilia can become part of an IO system. Phototactic steering is an important IO function that is specific to locomotion, and can be found in many metazoan larvae. The addition of this IO function does not require nervous control – in some sponge larvae, phototaxis is achieved by photosensitive ciliated cells that use light-controlled rudder-like cilia for phototactic steering without a need for nervous control [23,24]. So especially in cilia-based behavior, there can be significant IO and IC function without a nervous system.

In other cases, however, especially in bilaterians, the steering and modification of ciliary motion does come under nervous control. It has been proposed that the presence of a nervous system increases the efficiency of the control of cilia since a few sensors can control many effectors [6] (see also [4]). This allows the diversification of the repertoire of senses since a few dedicated sensory cells are sufficient to perform a specialized function.

Ciliary bands can be controlled directly by multifunctional sensory-motor cells, as during phototaxis in the larvae of the annelid *Platynereis dumerilii* (Figure 2) [25]. In the same larvae many different sensory-motor peptidergic neurons can control cilia to regulate the swimming depth of the larvae [26]. Large ciliomotor neurons can enable the control of all the cilia at once in a large body. In *Platynereis* larvae the simultaneous arrest of all cilia is triggered by one giant ciliomotor neuron (C Verasztó and GJ, unpublished). Via neural mechanisms, then, many effectors may be yoked to a smaller and specialized sensory apparatus.

**Figure 2.**
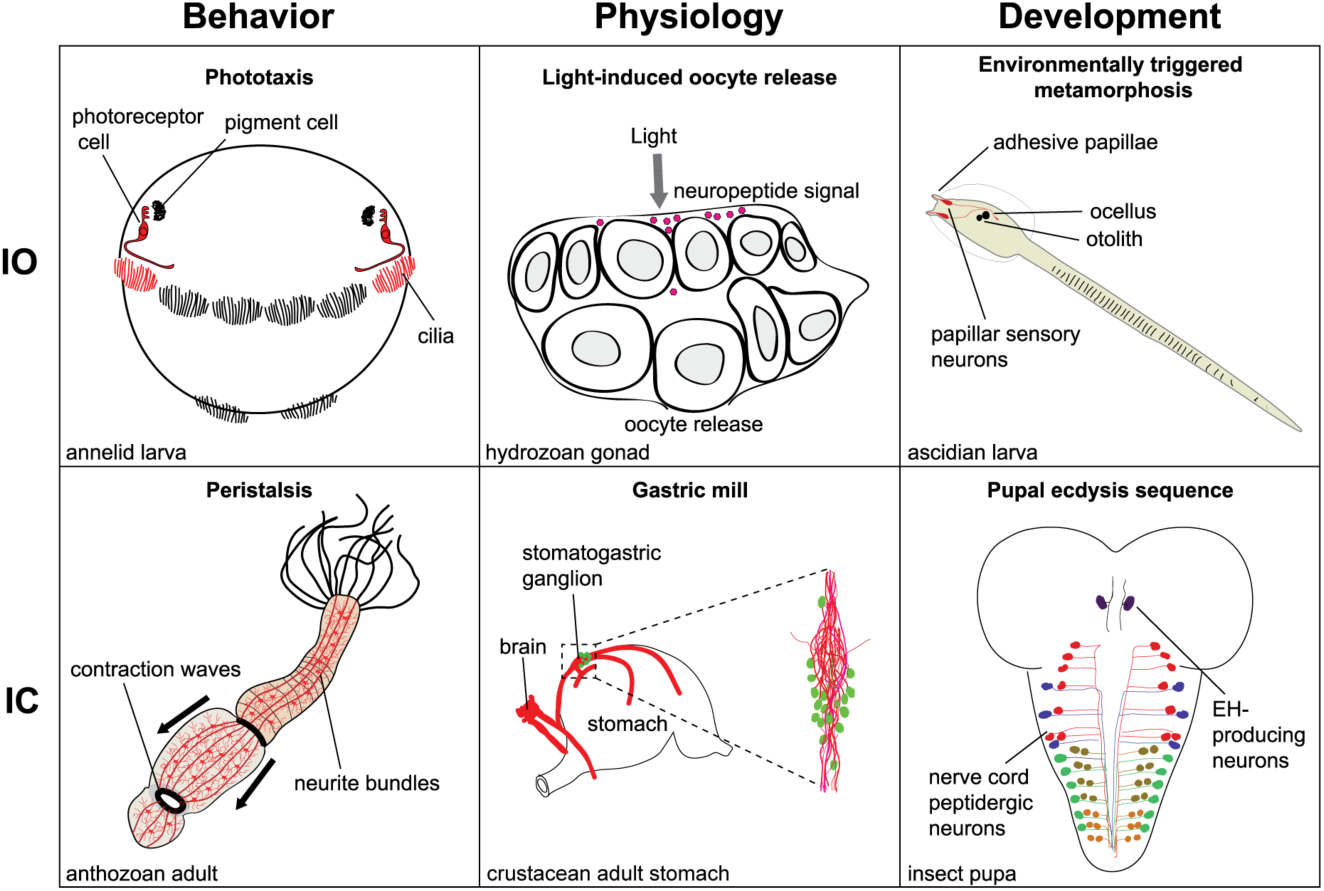
Examples of input-output and internal coordination systems for the control of behavior, physiology, and development.

Other cases show a similar role for efficiency enabled by division of labor. In the larvae of the mollusk *Helisoma* [sp], a few sensory neurons innervate the entire ciliary band and upregulate cilia beat frequency under hypoxic conditions [27]. Tosches and Arendt ([28]) have recently described neuroendocrine control of ciliary swimming in the larva of *Platynereis.* These larvae migrate from deeper to shallower water on a daily basis, and make use of a light-entrained neuroendocrine signal, melatonin, to control ciliary swimming. The use of a neuroendocrine system enables a damped response to changes in light input – the system does not respond to momentary changes in illumination. In the ctenophore *Pleurobrachia pileus,* prey capture triggers a complex and coordinated change in ciliary beating in the organism [29].

The addition that neurons make to ciliary systems, then, is on the IO side. Ciliary systems don’t require neural control for internal coordination. The role of nervous systems is to speed up cilia, slow them down, reverse and arrest them, in accordance with sensory information.

### 3.2 IC and IO systems for muscle-based behaviors

A second category of behavioral effector systems is muscle. As in the case of “neuron,” which we discussed above, giving a biologically reasonable definition of “muscle” is not a trivial matter. In the sense we use in this paper, a muscle is *an area of contractile cells with highly organized actin-myosin filaments.* Muscle in this sense includes myoepithelia (epithelia with contractile properties) in addition to smooth and striated muscle, but is absent from sponges and placozoa. Using this sense of “muscle,” together with the definition of “neuron” given above, an important generalization can be stated: *all organisms with nervous systems have muscle, and vice versa.* The only exceptions to this generalization are the myxozoan cnidarians, very reduced parasites, who have muscles but no nerves [30].

Some behaviors that involve contraction can be achieved without muscles or nerves; sponges use contractions in response to touch and to regulate the flow of water through their bodies and expel obstructions [31] [32] [33] [34]. These behaviors are fairly slow compared to those that can be achieved with muscle, however. Muscle-based effector systems require quite complex internal coordination to initiate and control events spread out across large multicellular structures. Such multicellular patterning of effector activity is where many important IC roles for nervous systems can be found.

Locomotion in pelagic animals involving jet propulsion and undulatory swimming are good examples of behaviors in which the animal body has to function as a coherent unit in order for muscle to work. In benthic organisms there is crawling, peristaltic movement, and “foot” movement in mollusks [35]. An important minimal example is a mixed form of locomotion seen in a hydrozoan larva *(Clava multicornis).* The larva propels itself along the substrate by ciliary motion, but steering is effected by lateral contractions of the body [36]. A slightly more complex organization is described for the acoel flatworm *Convoluta pulchra* [37]. Here cilia also drive the worm forward, but muscle allows the worm to change its shape and direction of movement as well as to position its mouth and ingest food. While ciliated surfaces require little neural control to act as a coordinated effector, the coordination of muscle surfaces is a more challenging problem – coordination does not come “for free,” as it does in ciliary metachronal waves.

Sessile organisms also have a range of uses for muscle, including sphincter movement, control of tentacles, gut peristaltic movements, and the pumping or release of gametes. In anthozoa, the larvae move by means of cilia, and the sessile adult uses muscle. The anemone *Nematostella* may illustrate an early role for muscle, as a replacement for ciliary methods for moving food within the organism. *Nematostella* has a long gut through which food is moved by peristaltic motion (Figure 2).

As discussed also in the case of ciliary motion, different models may be applicable to the initial laying down of a coordinated muscle-based effector and to later events by which finer control is added. Thus, when muscle coordination is in place, IO functions may become important. Simple examples of these roles include stopping a pattern of muscle contraction, speeding it up, switching between one pattern and another, and so on (e.g. [38]). In this case, muscles come under the partial control of specialized sensing devices and, complex neural structures that mediate between sensors and effectors can become prominent. These cases are highly familiar and we will not discuss them here.

### 3.3 IC and IO systems for gland-based behavior

Glandular systems are often neglected when discussing nervous system function or evolution. Such systems are very widespread and are essential in many animal groups for the normal execution of several behaviors, including predation, locomotion, and surface adhesion. During predation, several animals use special glands to catch or kill prey. These include the nematocyst and toxin-producing gland cells of cnidarians [39,40], the colloblasts of ctenophores [41], or the slime glands of onychophorans [42]. These structures are under nervous control [41,43] and sensory stimuli regulate secretion or discharge. For example, in cnidarians, toxin-gland and cnidocyte discharge is triggered by prey encounter [40], and is also influenced by light [44] (IO systems). In small interstitial marine invertebrates gland systems often contribute to locomotion by regulating surface adhesion, stopping, turning, or ciliary gliding. Interstitial animals, such as gastrotrichs, small annelids or flatworms can adhere to the substrate with the caudal parts of their bodies when disturbed. A specialized adhesive system, the duo-gland adhesive system, common among the gastrotrichs, secretes an adhesive substance from one type of gland and another substance from another gland that breaks the attachment [45]. Attachment can be triggered by mechanical disturbation (e.g. waves), and deattachment occures once the disturbance subsides, suggesting sensory/nervous control. In some cases adhesive glands are directly adjacent to a sensory neuron and nerves, suggesting nervous regulation (IO system) [46]. Adhesive glands can also contribute to turning and stopping, for example in *Monocelis,* a rapidly moving flatworm [45]. Another type of gland system contributes to ciliary gliding. Rhabdites are rod-shaped secretory products of some flatworms, nemerteans, gastrotrichs and annelids and are thought to provide a secreted layer of sticky mucus for ciliary gliding [47,48]. Mucus-secreting cells can also be under sensory control, as for example the mucus cells of ctenophores that are innervated by sensory-motor neurons (IO system) [49].

The above examples were more on the IO side, but several glandular secretion behaviors are under autonomous programs and represent IC systems, including tube-building by annelids using mucus-producing cells and sand grains [50,51]. The luminescent photocytes of ctenophores (that we discuss under gland systems) represent an interesting example showing both IC and IO aspects. Photocytes are innervated and are activated by mechanical stimulation (IO aspect) [52,53] but they are also electrically coupled, suggesting the spread of luminescence excitation among cells (IC aspect) [53].

## 4. IC and IO systems for the neural control of physiology

Although many discussions regard nervous systems as fundamentally concerned with control of behavior, these systems also have important roles in the control of physiology. This becomes self-evident when one considers the autonomic nervous system that coordinates functions like metabolism, internal clocks, digestion, heart rate and many other activities. Here, too, a distinction between IC and IO roles can be made, and often IC and IO functions are superimposed onto each other. As for behavior, also for physiological functions we can distinguish three types of effectors that the nervous system can influence, cilia, muscles, and glands. Some physiological processes require internal coordination which nervous systems make possible. Complex, muscle-driven physiological processes, such as peristaltic contractions to move the content of the gut or heartbeat, require IC systems to control them. For example, during defecation behavior in the fly, the hindgut and anal sphincter is driven by the sequential activation of motorneurons [54], representing an IC system. The motorneurons also receive sensory feedback from a mechanosensory neuron in the anus, adding an IO component to the circuit [54].

Gland-based systems can also use IC and IO mechanisms. Salivary gland cells in gastropod mollusks and the mouse are electrically coupled which allows the propagation of action potentials along the glandular epithelium, coordinating secretion from many cells [55,56]. This represents an example of an IC system of physiology in non-neural excitable cells. Saliva secretion can be induced by the nervous system [57], sometimes in a clear IO setting, as during the gustatory-salivary reflex, where taste inputs lead to saliva secretion [58].

Other aspects of the neural control of physiology also involve mixed IC and IO functions. A range of physiological functions are controlled by perception of light, especially by melatonin-based signaling systems. Melatonin signaling is very old, seen in cnidarians [59] and annelids [28] as well as chordates [60], and it can control both behavioral changes and several aspects of physiology, including sleep, appetite, and reproduction [60,61]. Many marine animals make use of moonlight to control the timing of reproduction [62]; corals, for example, spawn once each year in a way controlled by temperature, daily photoperiod and moonlight [63]. These are all IO functions; light is an external variable which must be tracked in some way. However, these IO functions influence the circadian or the circalunar clock that are intrinsically IC devices and are ultimately responsible for the periodicity of the physiological signals.

As during melatonin signaling, organisms often make use of neurosecretory mechanisms to determine a physiological response. In some hydrozoans, the release of oocytes is regulated by light-controlled neuroendocrine signaling [64], representing a clear IO case. Other neuroendocrine signaling systems have a mixed IO/IC character. Insulin-related peptides, produced by neurosecretory cells in flies and other metazoans, regulate glucose levels in the hemolymph, and lipid and carbohydrate storage. Dietary sugars, proteins, and lipids induce insulin release into the circulation by directly or indirectly affecting insulin-producing cells [65,66]. This and similar cases of internal homeostatic control by neuroendocrine mechanisms (e.g. diuresis, regulation of oxygen level in the blood) can be considered mixed IO/IC systems, with potentially deep evolutionary ancestry [67]. Homeostatic control in general requires a mechanism to sense internal states and an effector mechanism to change them (IO aspect). However, homeostatic control also has a strong IC character, due to the presence of negative feedback and that internal conditions are regulated.

The control of ciliated effector systems in physiology can also have IC and IO aspects. For example, the cilia of ependymal cells in the wall of the cerebral ventricles in vertebrates are regulated by neuron-derived melanin-concentrating hormone [68]. This system may be responsive to changes in glucose levels (IO aspect) and upregulate cerebrospinal fluid flow when glucose levels drop. Interestingly, as we have seen above for salivary gland cells, the adjacent ciliated ependymal cells are electrically coupled, allowing the coordination of activity (IC aspect) [69].

There are also several examples of the non-neural control of physiology by both IC and IO systems. The epidermal cilia of corals generate strong vortical flows that enhance the exchange of nutrients and dissolved gasses in the boundary layer. Whether the beating of cilia can be influenced by environmental cues (e.g. oxygen level) is not known in this system [70]. In other ciliated epithelia sensory stimuli are known to influence beating. Ciliated cells of human airway epithelial express sensory bitter taste receptors, and bitter compounds increase ciliary beat frequency by a cell-autonomous mechanism [71].

## 5. IC and IO systems for the neural control of development

Another often-neglected but very important aspect of nervous system function is control of development, including metamorphosis, growth, molting, and sexual maturation. Again a distinction can be made between IC and IO functions, and often there is a close interaction between the two.

Developmental changes are often cued by external events (IO systems). The metamorphosis of marine invertebrate larvae is the clearest example. In many marine organisms, metamorphosis is triggered by environmental cues, which indicate a suitable site for settlement of a planktonic larval stage [72]. This pattern is seen across corals, annelids, mollusks, ascidians and others [73–75] [76] [77]. A swimming larva encounters an environmental cue that is processed neurally [75]. A neuroendocrine cascade then triggers metamorphosis [78]. Sponge larvae, which do not have neurons, also settle and trigger metamorphosis in a roughly similar way, and this developmental transition in sponges is potentially significant in the earliest history of nervous systems [79]. The neuroendocrine signaling during larval settlement and metamorphosis employs the same or homologous signaling molecules, including nitric oxide (NO) and Wamide neuropeptides [77,79,80], suggesting deep evolutionary conservation [81]. Marine larval metamorphosis is triggered by environmental cues and represents an IO system. Other life-cycle transitions are internally coordinated. Sexual maturation in vertebrates [82], or ecdysis in insects is regulated by complex hormonal or peptidergic signaling. For example, in *Drosophila,* the ecdysis sequence is under the control of a peptidergic signaling cascade, involving the stepwise activation of peptidergic neurons [83].

Growth and regeneration are also influenced by the nervous system [84,85]. For example, insulin-like peptides, conserved in most metazoans, including placozoans [86], have likely ancient IC roles in the regulation of growth, as well as physiology (see above) [85,87].

In sum, though control of development is the least familiar role of nervous systems from a cognitive science point of view, it may have been especially important in early nervous system evolution. Simple marine organisms often exhibit dramatic changes between different modes of living and their accompanying morphologies. The triggering and coordination of metamorphosis for many animals is a crucially important life-cycle transition that places significant demands on control systems.

## 6. Possible Historical Sequences

We have discussed six categories. Each combines an *explanatory model* with a *function of nervous systems.* What sort of historical organization of these options might exist? One possibility is that some single one of our six options was the first or the most important factor in early nervous system evolution. Claims of this kind have been made, or in some cases implied, in a number of earlier discussions. For example, Jékely [6], described a historical sequence in which behavior (especially locomotion) is central to early nervous system evolution, and an IO pattern of explanation is applied. Jékely’s hypothesis is a modern version of scenarios sketched also by Parker [3] and Mackie [4] [5], with a focus on ciliary locomotion. In Jékely’s model, precursors of nervous systems arose to improve control of ciliary locomotion by means of division of labor and economies of scale. Consider, for example, the non-neural control of swimming in a sponge larva. Here sensory mechanisms influence the activity of cilia on the same cell, thereby steering the whole larva. This way of connecting sensory and motor capacities is notably inefficient, as every motor component needs its own sensor. It would be more efficient for a small number of sensory cells to control a large bank of motor devices, and this is what the advent of neurons makes possible. So one plausible account of the origin of nervous systems focuses on the efficient control of locomotion by sensory mechanisms in a ciliated swimming stage of an early metazoan; this is an *IO behavior* hypothesis.

In contrast, other hypotheses stress the importance of the control of muscle-based movement. The primacy of the nervous control of muscles in evolution is supported by the observation that muscle cells and nerve cells, bar one exception, always co-occur across animal diversity. Keijzer, van Duijn and Lyon [10] recently offered an IC based proposal where muscle-based behavior is held central. Their “skin brain thesis” conjectures that contractile and excitable epithelia provided a basic contractile organization that became more complex when neurons added long-distance projections. The proposal stresses that muscle contraction requires whole-body coordination and early nervous systems organized this coordination. For example, they introduce the concept of a *Pantin surface,* defined as the total contractile surface (or volume) that an animal has available for motility. Useful motility requires specific and stereotypical patterns of contraction and extension across this surface. New kinds of cell-to-cell interactions became important and nervous systems arose, on this view, to coordinate the micro-actions of cells into the macro-actions of whole organisms. In this view, the initial role of external sensors is comparatively minimal compared to IC. However, a control structure for contraction-based motility can have acted as a scaffold for the subsequent evolution of large-scale external sensors [88].

Both these hypotheses, and others like them, remain difficult to test at present, especially as extant organisms are highly evolved compared to the proposed basic configurations. For example, the well-studied hydromedusa *Aglantha digitale* exhibits both excitable epithelia and nerve nets. However, their interaction cannot be seen as primitive as this organism’s nervous system must have undergone a major evolutionary overhaul to accommodate and integrate two giant axons used for a fast escape response [89]. The general message remains that all extant ‘primitive organisms’ have a long evolutionary history since the first nervous systems arose and cannot be without further evidence be taken as representative for any primitive condition.

Single-factor hypotheses such as these represent one class of possibilities. According to hypotheses of this kind, the other roles for nervous systems were added later. At the other extreme, it is possible that *all* of our six options were important from very early days. In elementary form, all of these functions are seen in simple extant marine animals. Furthermore, examples of all six of our categories are seen, to varying degrees, in marine animals that *lack* nervous systems. This shows the importance, in principle, of all six of these forms of control to simple marine organisms. Sponge metamorphosis is controlled by chemosensing of the substrate (IO, development), and their larvae exhibit ciliary locomotion controlled by photoreception (IO, behavior); contractile motions of *Trichoplax* are a form an internally coordinated behavior (IC, behavior), influenced by sensory cues from food (IO, behavior) [90] that may also influence digestive enzyme secretion (IO, physiology). *Trichoplax* also has insulin, a peptide that regulates growth and physiology [86] (IC, physiology/development).

Given that all six of the functions that neurons can play are seen in simple form in-animals without nervous systems, it is plausible that all six of the roles we have discussed evolved in parallel from the beginning. There is, at least, quite a strong argument here against the exclusive importance of any of the six. Deep comparative studies have the potential to clarify the origin of some of the early roles nervous systems played. For example, both insulin-like peptides (IC role in physiology and development) and Wamides (IO role in triggering metamorphosis) are ancient molecules with broadly conserved functions. Opsins (IO roles in behavior and physiology) are also old, present in ctenophores, cnidarians and bilaterians.

However, in different contexts and at different evolutionary stages, some of our six options may have become more important than others. An important role may be played by body size, for example. *Ceteris paribus,* larger bodies will present more challenges for internal coordination in the control of behavior. If the first nervous system arose in a small animal, especially if it first appeared in the larval stage [91], this will probably reduce the need for IC functions in behavior. The possible evolutionary sequence due to Jékely that was outlined above, in which an IO-behavior function is central, is based on the assumption of a small organism in which the demands of internal coordination on behavior are not great. This raises the possibility, also, that early nervous systems may have played somewhat different roles at different stages in the life cycle of a single organism. A small, motile larva faced IO problems; a larger adult, perhaps drifting or sessile, faces IC problems. In many extant animals the larval stage uses cilia for motion while later developmental stages make use of muscle for behavior – this combination is seen in all animals that have a dispersing ciliated larval stage (e.g. annelids, mollusks, cephalochordates) [92].

## Discussion

The primary aim of this paper has been to chart the space of options for early neural evolution, and also to highlight a number of evolutionary possibilities that are often neglected. We organized the options with a three-way distinction between the functions nervous systems can play, and a two-way distinction between explanatory models. The resulting space of options is represented in Figure 2 and Table 1.

**Table 1.** The ‘explanatory model’ and ‘function’ matrix of the origin of nervous systems. The entries in the cells give examples, in simple organisms, of each of the six roles that nervous systems can play.

| *function*<br>*explanatory models* | neural control of<br>behavior | neural control of<br>physiology | neural control of<br>development |
| --- | --- | --- | --- |
| **input-output** | phototaxis;<br>chemotaxis; toxin<br>gland discharge | light-controlled<br>reproduction; salivary<br>reflex | environmentally-<br>triggered<br>metamorphosis |
| **internal coordination** | jet propulsion;<br>undulatory<br>swimming | gut peristalsis; diuresis<br>control; neuroendocrine<br>control of metabolism | neuroendocrine control<br>of growth; ecdysis |

In our discussion we mentioned cases where one explanatory model or the other, IO or IC, is paramount or more conspicuous and discussed examples where IC and IO functions are combined in a single form of activity. Most nervous system functions require a combination of internal coordination *and* the matching of acts with the state of the environment, tracked through the senses. The same may apply to behaviors that were important in early neural evolution. One possibility, which may be common, is a situation that features a “default” behavior produced by an IC system, along with an IO system that overrides or modifies it. The swimming-beat contraction in scyphozoan medusae is a good illustration of a default behavior in those organisms, and a substantial portion of the neural activity in a jellyfish goes into maintaining this rhythmic behavior [93,94]. Against that IC background, the jellyfish can also modify its behavior according to conditions it senses in its environment [38]. Similarly, ctenophores generally maintain a default pattern ciliary motion but reverse this motion when they touch prey [29].

Another way IC and IO functions can be combined is for a number of internally coordinated motor programs to be chosen, with none as default, according to the sensing of external events. Yet another possibility is a more seamless integration of the two kinds of function. One intriguing possibility here derives from the systematic feedback generated by an organism’s own movement that would link IC to IO functions in a more direct way [88] and which links up with current developments in embodied cognition (e.g. [95]). A more integrated view of this kind is perhaps the most accurate way to think about present-day human behavior. Still, even if this possibility is assumed, we maintain that the distinction between IC and IO functions is theoretically important; these are two fundamentally different tasks that must be handled by an organism, whether they are separable with respect to mechanisms or not.

We do not deny these complications, and the coarse-grained character of our central distinctions. However, we maintain that workers in this field do tend to slip into one conceptual framework or the other – often an IO framework – and see neural evolution through that lens. We think that the evidence available at this point suggests an important role for both kinds of control system in early neural evolution.

So although the main purpose of this paper is charting the option space itself, we think that existing evidence points towards some cautious conclusions about the importance of these options in possible historical scenarios. In particular, single-factor explanations are made unlikely by the existence of precursors in non-neural animals of all six of the functions for nervous systems we have distinguished. Considerable uncertainty remains about the phylogenetic relationships between sponges, ctenophores, placozoa, and other non-bilaterian animals [96–99]. These relationships can be expected to be important to hypotheses about early neural evolution [98,100–103].

While the actual historical scenario – or scenarios – for the evolution of the first nervous systems is beyond us for the foreseeable future, we do think that the six options that we sketched do provide a general layout of the major constraints that operated on this historical event, or events. Our aim is not to defend one of these options at the expense of another, but to stress their relevance as ways to focus on different neural functions that play complementary roles in understanding how and why nervous systems first arose, and subsequently evolved into the wide variety of these systems seen today.

## Acknowledgments

We thank Katrine Worsaae for advice on gland systems. We thank Eörs Szathmáry for informing GJ about the utility of option spaces as seen in the Parmenides Eidos system (https://www.parmenides-foundation.org/application/parmenides-eidos/).

## Authors’ Contributions

All authors contributed to the conception, design, and drafting of the article.

## References

1. Braitenberg, V. 1986 Vehicles: Experiments in synthetic psychology.

2. Sherrington, C. 1906 The Integrative Action of the Nervous System. Yale University Press.

3. Parker, G. H. 1919 The elementary nervous system. Philadelphia: Lippincott.

4. Mackie, G. 1970 Neuroid conduction and evolution of conducting tissues. Q Rev Biol 45, 319–332.

5. Mackie, G. O. 1990 The elementary nervous system revisited. Am Zool 30, 907–920. (doi:10.2307/3883448)

6. Jékely, G. 2011 Origin and early evolution of neural circuits for the control of ciliary locomotion. P R Soc B 278, 914–922. (doi:10.1098/rspb.2010.2027)

7. Pantin, C. 1956 The origin of the nervous system. Pubblicazioni della Stazione zoologica di Napoli.

8. Passano, L. M. 1963 Primitive nervous systems. P Natl Acad Sci Usa 50, 306–313.

9. de Ceccatty, M. P. 1974 The origin of the integrative systems: a change in view derived from research on coelenterates and sponges. Perspect. Biol. Med. 17, 379–390.

10. Keijzer, F., van Duijn, M. & Lyon, P. 2013 What nervous systems do: early evolution, input-output, and the skin brain thesis. Adaptive Behavior, 21, 67–85. (doi:10.1177/1059712312465330)

11. Nielsen, C. 2005 Trochophora larvae: cell-lineages, ciliary bands and body regions. 2. Other groups and general discussion. J. EXP. ZOOL 304, 401–447. (doi:10.1002/jez.b.21050)

12. Nielsen, C. 2004 Trochophora larvae: cell-lineages, ciliary bands, and body regions. 1. Annelida and Mollusca. J. EXP. ZOOL 302, 35–68. (doi:10.1002/jez.b.20001)

13. Lacalli, T. C. & Gilmour, T. 2001 Locomotory and feeding effectors of the tornaria larva of Balanoglossus biminiensis. Acta Zoologica 82: 117–126.

14. Tamm, S. L. 1984 Mechanical synchronization of ciliary beating within comb plates of ctenophores. J Exp Biol 113, 401–408.

15. Guirao, B. & Joanny, J.-F. 2007 Spontaneous creation of macroscopic flow and metachronal waves in an array of cilia. Biophys J 92, 1900–1917. (doi:10.1529/biophysj.106.084897)

16. Brumley, D. R., Polin, M., Pedley, T. J. & Goldstein, R. E. 2012 Hydrodynamic synchronization and metachronal waves on the surface of the colonial alga Volvox carteri. Physical Review Letters 109, 268102.

17. Mitchell, B., Stubbs, J. L., Huisman, F., Taborek, P., Yu, C. & Kintner, C. 2009 The PCP pathway instructs the planar orientation of ciliated cells in the Xenopus larval skin. Curr Biol 19, 924–929. (doi:10.1016/j.cub.2009.04.018)

18. Momose, T., Derelle, R. & Houliston, E. 2008 A maternally localised Wnt ligand required for axial patterning in the cnidarian Clytia hemisphaerica. Development 135, 2105–2113. (doi:10.1242/dev.021543)

19. Momose, T., Kraus, Y. & Houliston, E. 2012 A conserved function for Strabismus in establishing planar cell polarity in the ciliated ectoderm during cnidarian larval development. Development 139, 4374–4382. (doi:10.1242/dev.084251)

20. Guirao, B. et al. 2010 Coupling between hydrodynamic forces and planar cell polarity orients mammalian motile cilia. Nat Cell Biol 12, 341–350. (doi:10.1038/ncb2040)

21. Adamska, M., Degnan, S. M., Green, K. M., Adamski, M., Craigie, A., Larroux, C. & Degnan, B. M. 2007 Wnt and TGF-beta Expression in the Sponge Amphimedon queenslandica and the Origin of Metazoan Embryonic Patterning. PLoS ONE 2, e1031. (doi:10.1371/journal.pone.0001031)

22. Lapébie, P., Borchiellini, C. & Houliston, E. 2011 Dissecting the PCP pathway: One or more pathways? Bioessays 33, 759–768. (doi:10.1002/bies.201100023)

23. Leys, S. P. & Degnan, B. M. 2001 Cytological basis of photoresponsive behavior in a sponge larva. Biol Bull 201, 323–338.

24. Maldonado, M., Durfort, M., McCarthy, D. A. & Young, C. M. 2003 The cellular basis of photobehavior in the tufted parenchymella larva of demosponges. Mar Biol 143, 427–441. (doi:10.1007/s00227-003-1100-1)

25. Jékely, G., Colombelli, J., Hausen, H., Guy, K., Stelzer, E. H. K., Nédélec, F. & Arendt, D. 2008 Mechanism of phototaxis in marine zooplankton. Nature 456, 395–399. (doi:10.1038/nature07590)

26. Conzelmann, M., Offenburger, S.-L., Asadulina, A., Keller, T., Münch, T. A. & Jékely, G. 2011 Neuropeptides regulate swimming depth of Platynereis larvae. Proc Natl Acad Sci USA 108, E1174–83. (doi:10.1073/pnas.1109085108)

27. Kuang, S., Doran, S. A., Wilson, R. J. A., Goss, G. G. & Goldberg, J. I. 2002 Serotonergic sensory-motor neurons mediate a behavioral response to hypoxia in pond snail embryos. J Neurobiol 52, 73–83. (doi:10.1002/neu.10071)

28. Tosches, M. A., Bucher, D., Vopalensky, P. & Arendt, D. 2014 Melatonin Signaling Controls Circadian Swimming Behavior in Marine Zooplankton. Cell 159, 46–57. (doi:10.1016/j.cell.2014.07.042)

29. Tamm, S. L. & Moss, A. G. 1985 Unilateral ciliary reversal and motor responses during prey capture by the ctenophore Pleurobrachia. J Exp Biol 114, 443–461.

30. Kent, M. L. & Andree, K. B. 2001 Recent advances in our knowledge of the Myxozoa. Journal of Eukaryotic Microbiology 48: 395–413. (doi:10.1111/j.1550-7408.2001.tb00173.x)

31. Elliott, G. R. D. & Leys, S. P. 2007 Coordinated contractions effectively expel water from the aquiferous system of a freshwater sponge. J Exp Biol 210, 3736–3748. (doi:10.1242/jeb.003392)

32. Leys, S. P., Mackie, G. O. & Meech, R. W. 1999 Impulse conduction in a sponge. J Exp Biol 202, 1139–1150.

33. Nickel, M. 2004 Kinetics and rhythm of body contractions in the sponge Tethya wilhelma (Porifera: Demospongiae). J Exp Biol 207, 4515–4524. (doi:10.1242/jeb.01289)

34. Renard, E., Vacelet, J., Gazave, E., Lapébie, P., Borchiellini, C. & Ereskovsky, A. V. 2009 Origin of the neuro-sensory system: new and expected insights from sponges. Integr Zool 4, 294–308. (doi:10.1111/j.1749-4877.2009.00167.x)

35. Trueman, E. R. In press. The Locomotion of Soft-Bodied Animals. American Elsevier Publishing Co. (1975).

36. Piraino, S., Zega, G., Di Benedetto, C., Leone, A., Dell’Anna, A., Pennati, R., Candia Carnevali, D., Schmid, V. & Reichert, H. 2011 Complex neural architecture in the diploblastic larva of Clava multicornis (Hydrozoa, Cnidaria). J Comp Neurol 519, 1931–1951. (doi:10.1002/cne.22614)

37. Tyler, S. & Rieger, R. M. 1999 Functional morphology of musculature in the acoelomate worm, Convoluta pulchra (Plathelminthes). Zoomorphology 119, 127–142. (doi:10.1007/s004350050087)

38. Bailey, K. M. & Batty, R. S. 1983 A laboratory study of predation by Aurelia aurita on larval herring (Clupea harengus): experimental observations compared with model predictions. Mar Biol 72, 295–301. (doi:10.1007/BF00396835)

39. Beckmann, A. et al. 2015 A fast recoiling silk-like elastomer facilitates nanosecond nematocyst discharge. BMC Biol 13, 3. (doi:10.1186/s12915-014-0113-1)

40. Moran, Y., Genikhovich, G., Gordon, D., Wienkoop, S., Zenkert, C., Ozbek, S., Technau, U. & Gurevitz, M. 2012 Neurotoxin localization to ectodermal gland cells uncovers an alternative mechanism of venom delivery in sea anemones. Proc Biol Sci 279, 1351–1358. (doi:10.1098/rspb.2011.1731)

41. Franc, J. M. 1978 Organization and function of ctenophore colloblasts: an ultrastructural study. Biol Bull 155, 527–541.

42. Haritos, V. S., Niranjane, A., Weisman, S., Trueman, H. E., Sriskantha, A. & Sutherland, T. D. 2010 Harnessing disorder: onychophorans use highly unstructured proteins, not silks, for prey capture. Proc Biol Sci 277, 3255–3263. (doi:10.1098/rspb.2010.0604)

43. Westfall, J. A. 1970 The nematocyte complex in a hydromedusan, Gonionemus vertens. Z Zellforsch Mik Ana 110, 457–470. (doi:10.1007/BF00330098)

44. Plachetzki, D. C., Fong, C. R. & Oakley, T. H. 2012 Cnidocyte discharge is regulated by light and opsin-mediated phototransduction. BMC Biol 10, 17. (doi:10.1186/1741-7007-10-17)

45. Martin, G. G. 1978 The duo-gland adhesive system of the archiannelids Protodrilus and Saccocirrus and the turbellarian Monocelis. Zoomorphologie 91, 63–75. (doi:10.1007/BF00994154)

46. Tyler, S., Melanson, L. A. & Rieger, R. M. 1980 Adhesive organs of the gastrotricha. Zoomorphologie 95, 17–26. (doi:10.1007/BF01342231)

47. Martin, G. G. 1978 Ciliary gliding in lower invertebrates. Zoomorphologie 91, 249–261. (doi:10.1007/BF00999814)

48. Martin, G. G. 1978 A new function of rhabdites: Mucus production for ciliary gliding. Zoomorphologie 91, 235–248. (doi:10.1007/BF00999813)

49. Hernandez-Nicaise, M.-L. 1974 Ultrastructural evidence for a sensory-motor neuron in Ctenophora. Tissue & cell 6, 43–47. (doi:10.1016/0040-8166(74)90021-4)

50. Dorsett, D. A. 1961 The behaviour of *Polydora ciliata* (Johnst.). Tube-building and burrowing. J Mar Biol Assoc Uk 41, 577–590. (doi:10.1017/S0025315400016167)

51. Ziegelmeier, E. 1969 Neue Untersuchungen über die Wohnröhren-Bauweise vonLanice conchilega (Polychaeta, Sedentaria). Helgolander Wiss. Meeresunters 19, 216–229. (doi:10.1007/BF01625607)

52. Chang, J. J. 1954 Analysis of the luminescent response of the ctenophore, Mnemiopsis leidyi, to stimulation. J Cell Physiol 44, 365–394.

53. Anctil, D. M. 1985 Ultrastructure of the luminescent system of the ctenophore Mnemiopsis leidyi. Cell Tissue Res 242, 333–340. (doi:10.1007/BF00214545)

54. Zhan, W., Yan, Z., Li, B., Jan, L.Y. & Jan, Y.N. 2014 Identification of motor neurons and a mechanosensitive sensory neuron in the defecation circuitry of *Drosophila* larvae. eLife 3. (doi:10.7554/eLife.03293)

55. Kater, S. B. & Galvin, N. J. 1978 Physiological and morphological evidence for coupling in mouse salivary gland acinar cells. J Cell Biol 79, 20–26.

56. Kater, S. B., Rued, J. R. & Murphy, A. D. 1978 Propagation of action potentials through electrotonic junctions in the salivary glands of the pulmonate mollusc, Helisoma trivolvis. J Exp Biol 72, 77–90.

57. Marshall, C. G. & Lent, C. M. 1988 Excitability and secretory activity in the salivary gland cells of jawed leeches (Hirudinea: Gnathobdellida). J Exp Biol 137, 89–105.

58. Kawamura, Y. & Yamamoto, T. 1978 Studies on neural mechanisms of the gustatory-salivary reflex in rabbits. The Journal of Physiology 285, 35–47. (doi:10.1113/jphysiol.1978.sp012555)

59. Peres, R., Reitzel, A. M., Passamaneck, Y., Afeche, S., Cipolla-Neto, J., Marques, A. & Martindale, M. Q. 2014 Developmental and light-entrained expression of melatonin and its relationship to the circadian clock in the sea anemone Nematostella vectensis. EvoDevo 5, 26. (doi:10.1186/2041-9139-5-26)

60. Gandhi, A. V., Mosser, E. A., Oikonomou, G. & Prober, D. A. 2015 Melatonin Is Required for the Circadian Regulation of Sleep. Neuron 85, 1193–1199. (doi:10.1016/j.neuron.2015.02.016)

61. Lima Cabello, E., Díaz Casado, M. E., Guerrero, J. A., Otalora, B. B., Escames, G., López, L. C., Reiter, R. J. & Acuña Castroviejo, D. 2014 A review of the melatonin functions in zebrafish physiology. J. Pineal Res. 57, 1–9. (doi:10.1111/jpi.12149)

62. Tessmar-Raible, K., Raible, F. & Arboleda, E. 2011 Another place, another timer: Marine species and the rhythms of life. Bioessays 33, 165–72. (doi:10.1002/bies.201000096)

63. Richmond, R. H. & Hunter, C. L. 1990 Reproduction and recruitment of corals: Comparisons among the Caribbean, the Tropical Pacific, and the Red Sea. Marine Ecology Progress Series 60, 185–203.

64. Takeda, N., Nakajima, Y., Koizumi, O., Fujisawa, T., Takahashi, T., Matsumoto, M. & Deguchi, R. 2013 Neuropeptides trigger oocyte maturation and subsequent spawning in the hydrozoan jellyfish Cytaeis uchidae. Mol Reprod Dev 80, 223–232. (doi:10.1002/mrd.22154)

65. Broughton, S. J. et al. 2005 Longer lifespan, altered metabolism, and stress resistance in Drosophila from ablation of cells making insulin-like ligands. P Natl Acad Sci Usa 102, 3105–3110. (doi:10.1073/pnas.0405775102)

66. Brankatschk, M., Dunst, S., Nemetschke, L. & Eaton, S. 2014 Delivery of circulating lipoproteins to specific neurons in the Drosophila brain regulates systemic insulin signaling. Elife 3. (doi:10.7554/eLife.02862)

67. Tarrant, A. M. 2005 Endocrine-like Signaling in Cnidarians: Current Understanding and Implications for Ecophysiology. Am Zool 45, 201–214. (doi:10.1093/icb/45.1.201)

68. Conductier, G. et al. 2013 Melanin-concentrating hormone regulates beat frequency of ependymal cilia and ventricular volume. Nat Neurosci 16, 845–847. (doi:doi:10.1038/nn.3401)

69. Jarvis, C. R. & Andrew, R. D. 1988 Correlated electrophysiology and morphology of the ependyma in rat hypothalamus. Journal of Neuroscience 8, 3691–3702.

70. Shapiro, O. H., Fernandez, V. I., Garren, M., Guasto, J. S., Debaillon-Vesque, F. P., Kramarsky-Winter, E., Vardi, A. & Stocker, R. 2014 Vortical ciliary flows actively enhance mass transport in reef corals. Proc Natl Acad Sci USA 111, 13391–13396. (doi:10.1073/pnas.1323094111)

71. Shah, A., Ben-Shahar, Y., Moninger, T., Kline, J. & Welsh, M. 2009 Motile Cilia of Human Airway Epithelia Are Chemosensory. Science 325, 1131. (doi:10.1126/science.1173869)

72. Shikuma, N. J., Pilhofer, M., Weiss, G. L., Hadfield, M. G., Jensen, G. J. & Newman, D. K. 2014 Marine tubeworm metamorphosis induced by arrays of bacterial phage tail-like structures. Science 343, 529–533. (doi:10.1126/science.1246794)

73. Hadfield, M. G. & Paul, V. J. 2001 Natural chemical cues for settlement and metamorphosis of marine invertebrate larvae. Marine chemical ecology, 431–461.

74. Williams, E. & Degnan, S. 2009 Carry-over effect of larval settlement cue on postlarval gene expression in the marine gastropod Haliotis asinina. Molecular Ecology 18, 4434–4449. (doi:10.1111/j.1365-294X.2009.04371.x)

75. Hadfield, M. G., Meleshkevitch, E. A. & Boudko, D. Y. 2000 The apical sensory organ of a gastropod veliger is a receptor for settlement cues. Biol Bull 198, 67–76.

76. Cloney, R. A. 1982 Ascidian larvae and the events of metamorphosis. Am Zool 22, 817–826. (doi:10.2307/3882686)

77. Whalan, S., Webster, N. S. & Negri, A. P. 2012 Crustose Coralline Algae and a Cnidarian Neuropeptide Trigger Larval Settlement in Two Coral Reef Sponges. PLoS ONE 7, e30386. (doi:10.1371/journal.pone.0030386)

78. Heyland, A. & Moroz, L. L. 2006 Signaling mechanisms underlying metamorphic transitions in animals. Integrative and Comparative Biology 46, 743–759. (doi:10.1093/icb/icl023)

79. Degnan, S. M. & Degnan, B. M. 2010 The initiation of metamorphosis as an ancient polyphenic trait and its role in metazoan life-cycle evolution. Philos TRoy Soc B 365, 641–651. (doi:10.1098/rstb.2009.0248)

80. Conzelmann, M., Williams, E. A., Tunaru, S., Randel, N., Shahidi, R., Asadulina, A., Berger, J., Offermanns, S. & Jékely, G. 2013 Conserved MIP receptor-ligand pair regulates Platynereis larval settlement. Proc Natl Acad Sci U S A 110, 8224–8229. (doi:10.1073/pnas.1220285110)

81. Schoofs, L. & Beets, I. 2013 Neuropeptides control life-phase transitions. Proc Natl Acad Sci USA 110, 7973–7974. (doi:10.1073/pnas.1305724110)

82. Messager, S. et al. 2005 Kisspeptin directly stimulates gonadotropin-releasing hormone release via G protein-coupled receptor 54. P Natl Acad Sci Usa 102, 1761–1766. (doi:10.1073/pnas.0409330102)

83. Kim, Y.-J., Zitňan, D., Galizia, C. G., Cho, K.-H. & Adams, M. E. 2006 A command chemical triggers an innate behavior by sequential activation of multiple peptidergic ensembles. Curr Biol 16, 1395–1407. (doi:10.1016/j.cub.2006.06.027)

84. Kreshchenko, N. D., Sedelnikov, Z., Sheiman, I. M., Reuter, M., Maule, A. G. & Gustafsson, M. K. S. 2007 Effects of neuropeptide F on regeneration in Girardia tigrina (Platyhelminthes). Cell Tissue Res 331, 739–750. (doi:10.1007/s00441-007-0519-y)

85. Rulifson, E. J., Kim, S. K. & Nusse, R. 2002 Ablation of insulin-producing neurons in flies: growth and diabetic phenotypes. Science 296, 1118–1120. (doi:10.1126/science.1070058)

86. Nikitin, M. 2014 Bioinformatic prediction of Trichoplax adhaerens regulatory peptides. Gen Comp Endocrinol 212, 145–155. (doi:10.1016/j.ygcen.2014.03.049)

87. Brogiolo, W., Stocker, H., Ikeya, T., Rintelen, F., Fernandez, R. & Hafen, E. 2001 An evolutionarily conserved function of the Drosophila insulin receptor and insulin-like peptides in growth control. Curr Biol 11, 213–221.

88. Keijzer, F. 2015 Moving and sensing without input and output: early nervous systems and the origins of the animal sensorimotor organization. Biol Philos 30, 311–331.

89. Mackie, G. 2004 Central neural circuitry in the jellyfish Aglantha - A model ‘simple nervous system’. Neurosignals 13, 5–19. (doi:10.1159/000076155)

90. Ueda, T., Koya, S. & Maruyama, Y. K. 1999 Dynamic patterns in the locomotion and feeding behaviors by the placozoan Trichoplax adhaerence. Biosystems 54, 65–70.

91. Nielsen, C. 2013 Life cycle evolution: was the eumetazoan ancestor a holopelagic, planktotrophic gastraea? BMC Evol Biol 13, 171. (doi:10.1186/1471-2148-13-171)

92. Stokes, M. 1997 Larval locomotion of the lancelet Branchiostoma floridae. J Exp Biol 200, 1661–1680.

93. Passano, L. M. 1965 Pacemakers and activity patterns in medusae: homage to Romanes. Am Zool 5, 465–481. (doi:10.2307/3881171)

94. Schwab, W. E. 1977 The ontogeny of swimming behavior in the scyphozoan, Aurelia aurita. I. Electrophysiological analysis. Biol Bull 152, 233–250.

95. O’Regan, J. K. & Noë, A. 2001 A sensorimotor account of vision and visual consciousness. The Behavioral and brain sciences 24, 939–73-discussion 973–1031.

96. Moroz, L. L. et al. 2014 The ctenophore genome and the evolutionary origins of neural systems. Nature 510, 109–114. (doi:10.1038/nature13400)

97. Ryan, J. F. et al. 2013 The Genome of the Ctenophore Mnemiopsis leidyi and Its Implications for Cell Type Evolution. Science 342, 1242592–1242592. (doi:10.1126/science.1242592)

98. Jékely, G., Paps, J. & Nielsen, C. 2015 The phylogenetic position of ctenophores and the origin (s) of nervous systems. EvoDevo 6, 1.(doi:10.1186/2041-9139-6-1)

99. Philippe, H. et al. 2009 Phylogenomics revives traditional views on deep animal relationships. Curr Biol 19, 706–712. (doi:10.1016/j.cub.2009.02.052)

100. Moroz, L. L. 2015 Convergent evolution of neural systems in ctenophores. Journal of Experimental Biology 218, 598–611. (doi:10.1242/jeb.110692)

101. Ryan, J. F. 2014 Did the ctenophore nervous system evolve independently? Zoology (Jena) 117, 225–226. (doi:10.1016/j.zool.2014.06.001)

102. Marlow, H. & Arendt, D. 2014 Evolution: ctenophore genomes and the origin of neurons. Curr Biol 24, R757–61. (doi:10.1016/j.cub.2014.06.057)

103. Dunn, C. W., Leys, S. P. & Haddock, S. H. D. 2015 The hidden biology of sponges and ctenophores. Trends Ecol Evol (Amst) 30, 282–291. (doi:10.1016/j.tree.2015.03.003)

